# Stable gene expression for normalisation and single-sample scoring

**DOI:** 10.1101/2020.05.04.077859

**Authors:** Dharmesh D. Bhuva, Joseph Cursons, Melissa J. Davis

## Abstract

**Background:** Transcriptomic signatures are useful in defining the molecular phenotypes of cells, tissues, and patient samples. Their most successful and widespread clinical application is the stratification of breast cancer patients into molecular (PAM50) subtypes. In most cases, gene expression signatures are developed using transcriptome-wide measurements, thus methods that match signatures to samples typically require a similar degree of measurements. The cost and relatively large amounts of fresh starting material required for whole-transcriptome sequencing has limited clinical applications, and accordingly thousands of existing gene signatures are unexplored in a clinical context.

**Results:** Genes in a molecular signature can provide information about molecular phenotypes and their underlying transcriptional programs from tissue samples, however determining the transcriptional state of these genes typically requires the measurement of all genes across multiple samples to allow for comparison. An efficient assay and scoring method should quantify the relative abundance of signature genes with a minimal number of additional measurements. We identified genes with stable expression across a range of abundances, and with a preserved relative ordering across large numbers (thousands) of samples, allowing signature scoring, and supporting general data normalisation for transcriptomic data. Based on singscore, we have developed a new method, *stingscore*, which quantifies and summarises relative expression levels of signature genes from individual samples through the inclusion of these “stably-expressed genes”.

**Conclusion:** We show that our proposed list of stable genes has better stability across cancer and normal tissue data than previously proposed stable or housekeeping genes. Additionally, we show that signature scores computed from whole-transcriptome data are comparable to those calculated using only values for signature genes and our panel of stable genes. This new approach to gene expression signature analysis may facilitate the development of panel-type tests for gene expression signatures, thus supporting clinical translation of the powerful insights gained from cancer transcriptomic studies.

## Introduction

Measurements of the transcriptome are often used to infer the molecular phenotype of a biological system. Assays that capture such measurements vary in throughput, accuracy, and cost (1). Real time quantitative PCR (RT-qPCR) is an accurate assay with low throughput, whereas RNA-seq is a high throughput assay that is relatively less accurate. Reduced costs coupled with accurate measurements make RT-qPCR a popular platform for clinical RNA-based tests. The Oncotype Dx® is a prognostic test that measures the abundance of 21 genes in breast cancer using RT-qPCR to compute scores that predict the risk of recurrence and guide chemotherapy (2).

NanoString’s nCounter® platform can provide moderate-throughput measurements of several hundred transcripts at a low cost, and this platform is used in the PAM50-based Prosigna® breast cancer prognostic gene signature assay (3). This test first classifies samples into PAM50 subtypes and then uses these estimates to predict a risk or recurrence score. PAM50 subtypes are higher order molecular phenotypes of patient tumours that capture information on their molecular state. Lower order phenotypes that probe the microenvironment can be used to assess tumour infiltrating lymphocytes and potentially guide immunotherapies (4). Likewise, phenotypes that assess pathway activity may be used to predict drug sensitivity and consequently guide therapy (5).

In research, these phenotypes are routinely assessed using transcriptomic gene signatures such as those available in the molecular signature database (MSigDB) (6,7), and insights provided by them have been vital in developing our understanding of cancer. Tens of thousands of gene signatures have been developed and used in a research context, but their clinical potential remains largely unexplored. This limitation is due to the need for transcriptome-wide measurements for most molecular phenotyping methods, which is not cost-effective and may not be feasible in a clinical setting. For instance, samples that are formalin fixed and paraffin embedded (FFPE) are not suitable for whole transcriptome RNA-seq. As a consequence, large and valuable collections of archival material cannot be used for RNA-seq based transcriptomics, but they may be amenable to alternative, low-to medium-throughput cost-effective methods like RT-qPCR or NanoString nCounter®. Supporting this approach, the testing of archival FFPE samples has shown targeted pathway methods to be the most reliable form of transcriptomic analysis for these samples (8). This highlights a need for systematic methods for translating whole transcriptome derived gene expression signatures to a reduced measurement space allowing the potential exploration of archival samples and supporting the development of further targeted clinical assays.

A core requirement across all platforms is the need for control genes to normalise across sample measurements. Sample-to-sample variation can arise either form biological or technical sources. The effects of some technical variation can be minimised by normalising measurements against controls; this variation can arise from differences in total starting material, enzymatic efficiencies and in transcriptional activity between tissues or cells (9). More complex forms of variation such as batch effects can be corrected with sophisticated approaches that make use of control genes (10). Experimental spike-ins such as External RNA Controls Consortium (ERCC) spike-ins (11) can be used as control genes, however they do not experience the same sample preparation steps as endogenous RNA and therefore may not be the best representative reference (12–14). Historically, *ACTB* and *GAPDH* have been used as endogenous controls in RT-qPCR experiments, however, numerous studies have shown them to be differentially expressed across tissues (15–18). Alternatively, *housekeeping genes* that are assumed to be invariant across tissues due to their involvement in core cellular processes can be used as endogenous controls (19), however they have also been shown to vary across different tissues (9,15,20,21).

The availability of large transcriptomic datasets spanning numerous biological conditions has encouraged data-driven identification of reference genes. Vandesompele et al. (9) introduced the geNorm algorithm to select reference genes for microarray data by iterative elimination of the least stable gene. Key ideas they introduced were the use of multiple genes for normalisation, and evaluation of gene stability with respect to a putative set of stable genes. Reference genes for the NanoString nCounter® pan-cancer panels were selected by applying geNorm to the genotype-tissue expression dataset (GTEx). Krasnov et al. (20) used pan-cancer and normal RNA-seq data from the cancer genome atlas (TCGA) to prioritise reference genes for RT-qPCR experiments on tumour samples. Genes were prioritised such that they were differentially expressed between tumour and normal samples, had low variation as measured by the standard deviation, were associated with clinical parameters and had a high expression. Genes with many mutations, isoforms and pseudogenes were penalised. All information was weighted and collated using heuristic scoring functions. Lin et al. (21) have identified stable genes for normalisation single cell RNA sequencing (scRNA-seq) data. Using gamma-Gaussian mixture models for gene expression, they decomposed the lower expression spectrum of each gene into a gamma distribution. Next, they prioritised genes with a smaller gamma component, lower variation at the higher end of the expression spectrum, a smaller proportion of zero counts, and lower differences between cell clusters. Their approach penalised genes with low expression across cell clusters.

Both data-driven approaches above defined stably expressed genes by computing multiple measures of stability and combining them into a single metric that can be used to prioritise stably expressed genes. A shared limitation of these studies was dependence on a single dataset. Meta-analysis approaches generally produce robust results and have been widely used in differential expression analysis. As such, using multiple datasets to identify stable genes would produce a more robust prioritisation. Additionally, Krasnov et al. (20) proposed a cancer-specific set of stable genes but only used data from tumours to define their list. Most cancer-specific analyses are initially performed on cell lines and later translated to patients or patient derived xenografts. As such, genes used to calibrate these datasets need to be stable in both tumours and other experimental models.

In this study, we aim to address these limitations and improve the process of prioritising stable genes. We compute a variety of stability measures across two diverse datasets and combine them using a meta-analysis method (22). A measure of outliers is included to ensure stability is maintained across as many samples as possible, and to allow usage of these genes in outlier-based analyses (23). The list of stably expressed genes we propose are comparable or better than other lists in terms of stability while possessing additional properties. Our list covers a wider range of expression values than previous studies and therefore may better capture variability towards the tails of the expression distribution. A novel property of rank preservation is observed whereby the relative ranks of stable genes are also preserved across samples. This additional information may provide better opportunities for some normalisation methods, and for rank-based analysis methods. Finally, we demonstrate how the list of stable genes along with information on their relative ranks may provide cost-effective opportunities to test for molecular signature enrichment. The lists we provide in this study may be used in a diverse set of applications, including, RT-qPCR, NanoString nCounter®, and RUV-based normalisation (10).

## Materials and methods

### Pre-processing datasets

Where count-level data were available, gene filtering was performed on log-transformed counts-per-million reads (logCPM) with subsequent trimmed mean of m-values (TMM) normalisation (24) and finally transformation to log-transformed reads per kilobase of transcript, per million reads (logRPKM). RNA-seq data from post-mortem samples were obtained through the Genotype-Tissue Expression consortium (GTEx). Some samples have undergone autolysis which results in poor RNA quality, therefore samples with autolysis scores greater than 1 were excluded from the analysis. Pan-cancer RNA-seq data from The Cancer Genome Atlas (TCGA) were obtained as Subread-processed output from GEO (GSE62944). Samples with the phrase “carcinoma” in their annotation were considered carcinomas and were used to derive our stable gene list. TCGA breast cancer data were downloaded and processed using an alternate pipeline described in a R/Bioconductor-based workflow (25). This data was used to assess the impact of processing pipelines on putative stable genes. Cancer cell line encyclopedia (CCLE) samples were classified as carcinomas similar to TCGA samples. CCLE data downloaded using the `PharmacoGx` R package (26) were used in this analysis. Finally, genes with a logRPKM, log-transformed transcripts per million (logTPM), logCPM, or log-transformed fragments per kilobase of transcript, per million reads (logFPKM) of less than 1 across more than half of the samples were filtered out from each dataset due to low abundance. Filtering to remove genes with low abundance was performed independently for carcinoma and non-carcinoma samples.

Sequencing quality control consortium (SEQC) RNA-seq data were obtained from the `seqc` R/Bioconductor package. RNA-seq measurements from Illumina instruments at the Australian Genome Research (AGR) centre processed using the RefSeq annotation were used in this study. RT-qPCR measurements from the PRIME-qPCR protocol were obtained from GEO (GSE56457).

### Computing metrics of variability

Four metrics of variability were computed for each gene within each dataset:

- The median absolute deviation (MAD) – a rank-based measure of variation. For gene expression measurements *X*_1_, *X*_2_, …, *X*_*n*_, where *X_i_*, is the expression of gene *i* across *m* samples, the MAD is calculated as 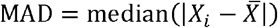 where 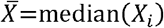.
- Shannon’s entropy – a measure of information content for a variable. The Shannon entropy is computed using the distribution of the data. We estimate the distribution by discretising gene expression measurements into bins of equal width using the `entropy` R package. The number of bins was computed as the square root of the number of samples and bins were defined using the expression distribution of the entire dataset. Shannon’s entropy for gene *i* is then computed as 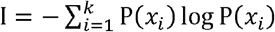 where *P*(*X*) is the probability mass function and *k* is the number of bins.
- The outlier sum statistic – a metric quantifying the presence of outliers. Outliers are defined as observations where transcript abundance is either greater than the sum of the third quartile and the interquartile range (IQR), or less than the difference of the first quartile and the interquartile range (*x* < *q*_0.25_ − IQR or *x* > *q*_0.75_ + IQR). The outlier sum statistic is then the sum of absolute value from median-centred outliers.
- F-statistic from a one-way ANOVA test on source tissue to assess group-wise differences.

R code used to compute these metrics and combine the results is available in additional file 3.

### Stability scores for gene sets

We repurposed singscore to compute gene set stability scores instead of enrichment scores. Genes were ranked based on their stability across all TCGA carcinoma samples and CCLE carcinoma-derived cell lines. We then created a single pseudo-sample with genes ranked on stability and computed uncentred scores for this pseudo-sample against gene sets using the R/Bioconductor package `singscore`. The resulting scores are each in the range of [0,1] with 1 indicating perfect gene set stability relative to all assessed genes. Gene sets were downloaded from MSigDB v5.2 (7). We discarded gene sets where fewer than 10 member genes had been assessed for stability in our study. The remaining gene sets were scored for stability using the approach described.

### Computing gene set scores using stable genes

The *stingscore* approach to scoring gene sets using stable genes is implemented in the R/Bioconductor package `singscore` (v1.8.0). Expression data can be ranked against stable genes by passing a set of stable genes to the `rankGenes()` function using the `stableGenes` argument. Passing the rank matrix to the `simpleScore()` function automatically invokes the *stingscore* implementation.

## Results

### Selecting stably expressed genes

In our study we have explored expression stability of genes relative to other genes. As such, variation of a gene across samples could be biologically significant, but it would be considered stable if it is less variable than other genes. In effect, like previous studies, we identified genes with a smaller dynamic range relative to other genes. Two cancer datasets representing different cancer models were used to identify stable genes: The Cancer Genome Atlas (TCGA) pan-cancer tumour data and Cancer Cell Line Encyclopedia (CCLE) cell line data (26,27). As the transcriptome of solid tumours are clearly distinct from those of liquid (haematological and lymphoid) malignancies, we have focused on identifying stable genes in solid tumours for this study. Thus, only carcinomas from TCGA and carcinoma-derived cell lines from the CCLE dataset were used to identify stable genes. Using multiple variability measurements across diverse datasets ensured a robust selection of stable genes.

For each gene, we calculated the median absolute deviation (MAD), Shannon’s entropy, the outlier sum statistic, and the F-statistic from a one-way ANOVA analysis - on either the tissue of the primary tumours or the tissue of origin for cell lines. The first three metrics quantify variability of gene expression while the last quantifies between group differences in abundance. Ideal stable genes would be invariant to tissue types. Hexbin plots in the background of Figure 1 show distributions of these measurements for TCGA carcinomas. These four quantities were measured for all genes across both datasets, resulting in eight measurements of variation.

**Figure 1.**
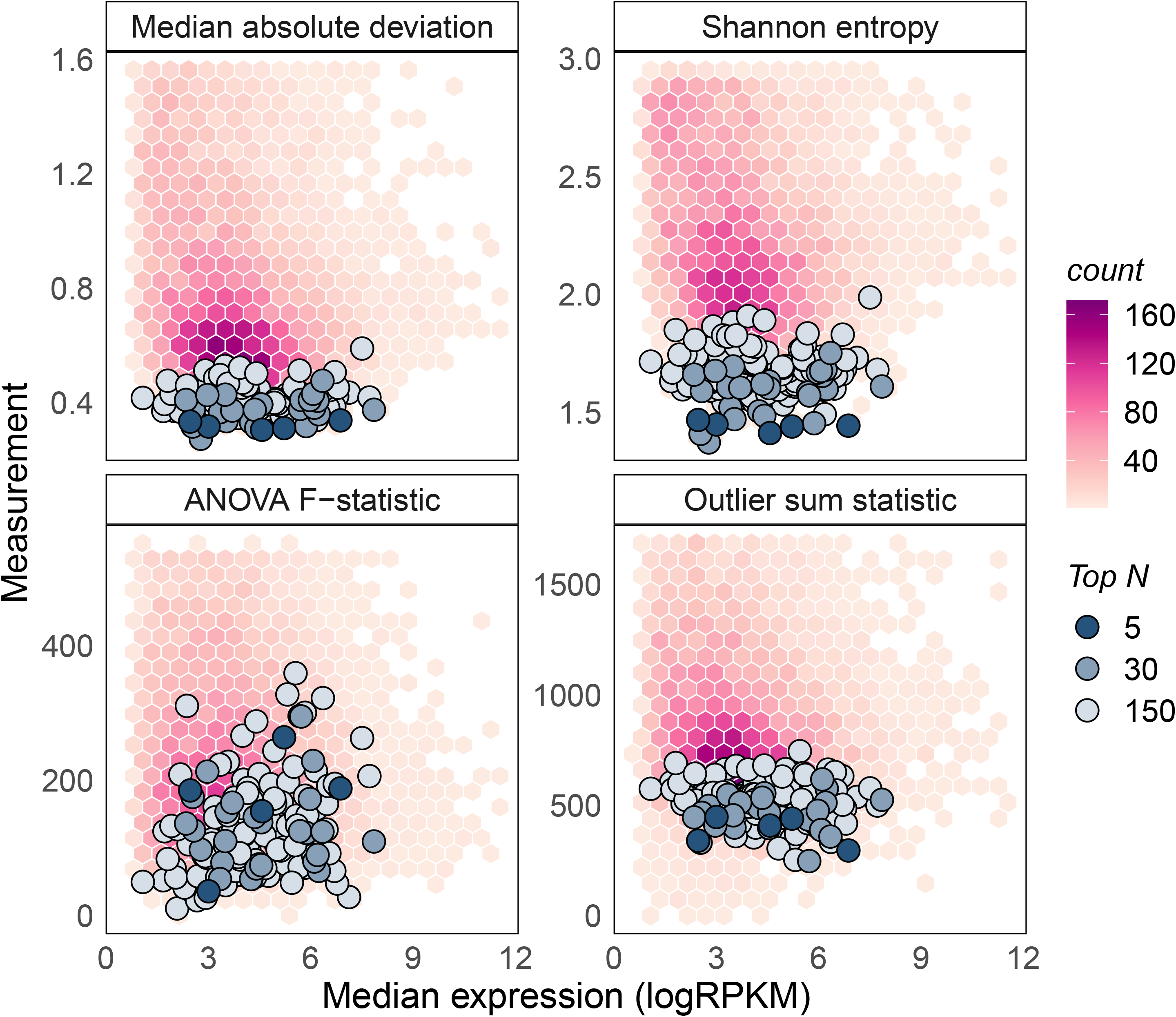
Stability metrics for stable genes are minimised on TCGA PanCancer carcinomas. The background distribution represents metrics for all genes. Top 5, 30 and 150 stable genes are overlayed on each plot. These genes have lower median absolute deviations, Shannon’s entropy and outlier sum statistics. They have relatively lower between group differences as signified by the F-statistic from the one-way ANOVA test on tissues. Stable genes tend to cover a wide range of the expression spectrum (widely distributed median logRPKMs).

Considering each of the eight variation metrics as separate stability analyses, we can use meta-analysis approaches to prioritise stable genes. We used the *product of ranks* rank aggregation method to combine stability information gained from the measurements (22). Briefly, genes were ranked in ascending order of each variability measurement independently resulting in eight *rankers*. These rankers were then combined using the product function to produce a product of ranks statistic for each gene. Genes missing in either dataset were discarded as their product of ranks could not be computed. A schematic of this approach is presented in the supplementary material (additional file 1: supplementary figure 1). The proposed list of stable genes can be accessed using the `getStableGenes()` function in the R/Bioconductor package `singscore` (v1.8.0). The top 5, 30 and 150 stable genes prioritised using this approach are overlayed on TCGA metric distributions in Figure 1. It is evident that the product of ranks approach prioritises genes that minimise most measures of variation. The F-statistic from the ANOVA analysis is higher for proposed stable genes. Variability measurements were plotted against the median expression value of each gene in Figure 1, demonstrating that our set of top stable genes covers a wide range of expression values.

### Comparison against other lists of stable genes

We used 14 independent datasets to assess the validity of our prioritisation of stable genes. These data are listed in Table 1 with details on the number of samples, groups and measurements, and the type of data or measurement. They are derived from tumour samples, cancer cell lines, normal tissue, and primary cell lines; with RNA-seq, proteomic and CAGE-seq measurements. Pre-processing was different for some of the data with RNA-seq and CAGE-seq measurements summarised as either TPM, CPM or FPKM/RPKM.

**Table 1.**
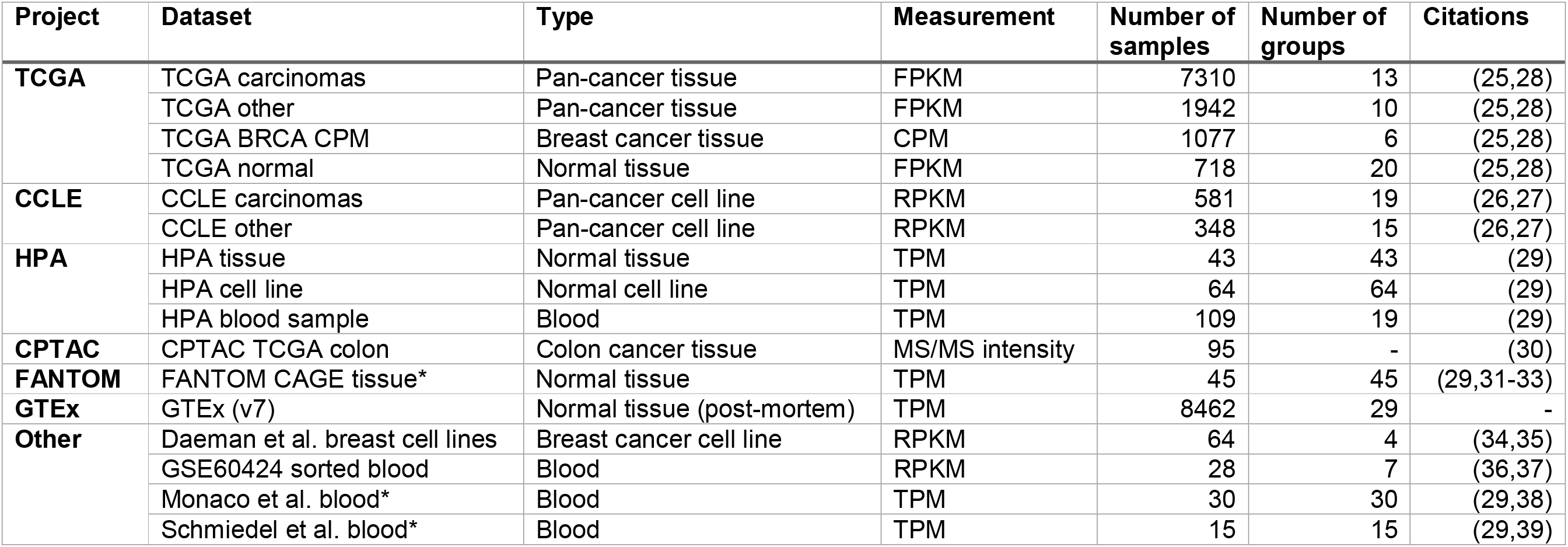
Datasets used for identifying and validating stably expressed genes (processed versions of datasets marked by asterisk were downloaded from the human protein atlas www.proteinatlas.org)

| Project | Dataset | Type | Measurement | Number of samples | Number of groups | Citations |
| --- | --- | --- | --- | --- | --- | --- |
| **TCGA** | TCGA carcinomas | Pan-cancer tissue | FPKM | 7310 | 13 | (25,28) |
|  | TCGA other | Pan-cancer tissue | FPKM | 1942 | 10 | (25,28) |
|  | TCGA BRCA CPM | Breast cancer tissue | CPM | 1077 | 6 | (25,28) |
|  | TCGA normal | Normal tissue | FPKM | 718 | 20 | (25,28) |
| **CCLE** | CCLE carcinomas | Pan-cancer cell line | RPKM | 581 | 19 | (26,27) |
|  | CCLE other | Pan-cancer cell line | RPKM | 348 | 15 | (26,27) |
| **HPA** | HPA tissue | Normal tissue | TPM | 43 | 43 | (29) |
|  | HPA cell line | Normal cell line | TPM | 64 | 64 | (29) |
|  | HPA blood sample | Blood | TPM | 109 | 19 | (29) |
| **CPTAC** | CPTAC TCGA colon | Colon cancer tissue | MS/MS intensity | 95 | - | (30) |
| **FANTOM** | FANTOM CAGE tissue* | Normal tissue | TPM | 45 | 45 | (29,31-33) |
| **GTEx** | GTEx (v7) | Normal tissue (post-mortem) | TPM | 8462 | 29 | - |
| **Other** | Daeman et al. breast cell lines | Breast cancer cell line | RPKM | 64 | 4 | (34,35) |
|  | GSE60424 sorted blood | Blood | RPKM | 28 | 7 | (36,37) |
|  | Monaco et al. blood* | Blood | TPM | 30 | 30 | (29,38) |
|  | Schmiedel et al. blood* | Blood | TPM | 15 | 15 | (29,39) |

We evaluated our prioritisation of stable genes on these datasets and simultaneously compared it against other stable gene lists or prioritisations. Specifically, we compared our rankings against those from the ordered lists of Lin et al. (21) and Krasnov et al. (20). We also compared stability against the set of stable genes used for normalisation in the NanoString nCounter® PanCancer pathways, PanCancer immune profiling and PanCancer progression gene expression panels. These panels have 40, 40 and 30 stable genes, respectively. To enable comparison of both discrete and prioritised lists of stable genes, we computed M-values proposed in the geNorm method (9). M-values are computed to capture the variability of each gene relative to every other gene in a putative set of stable genes. Thus, adding a gene to the set of stable genes will alter the M-value for all other genes in the set. We computed the median M-value and the interquartile range of M-values for a given set of stable genes. These values were computed for sets of size 5 to 150 for prioritisation lists such as ours. These quantities are shown in Figure 2 where the median M-value is represented as either a point or a line, and the interquartile range as error bars/bands. Lower M-values indicate better stability.

**Figure 2.**
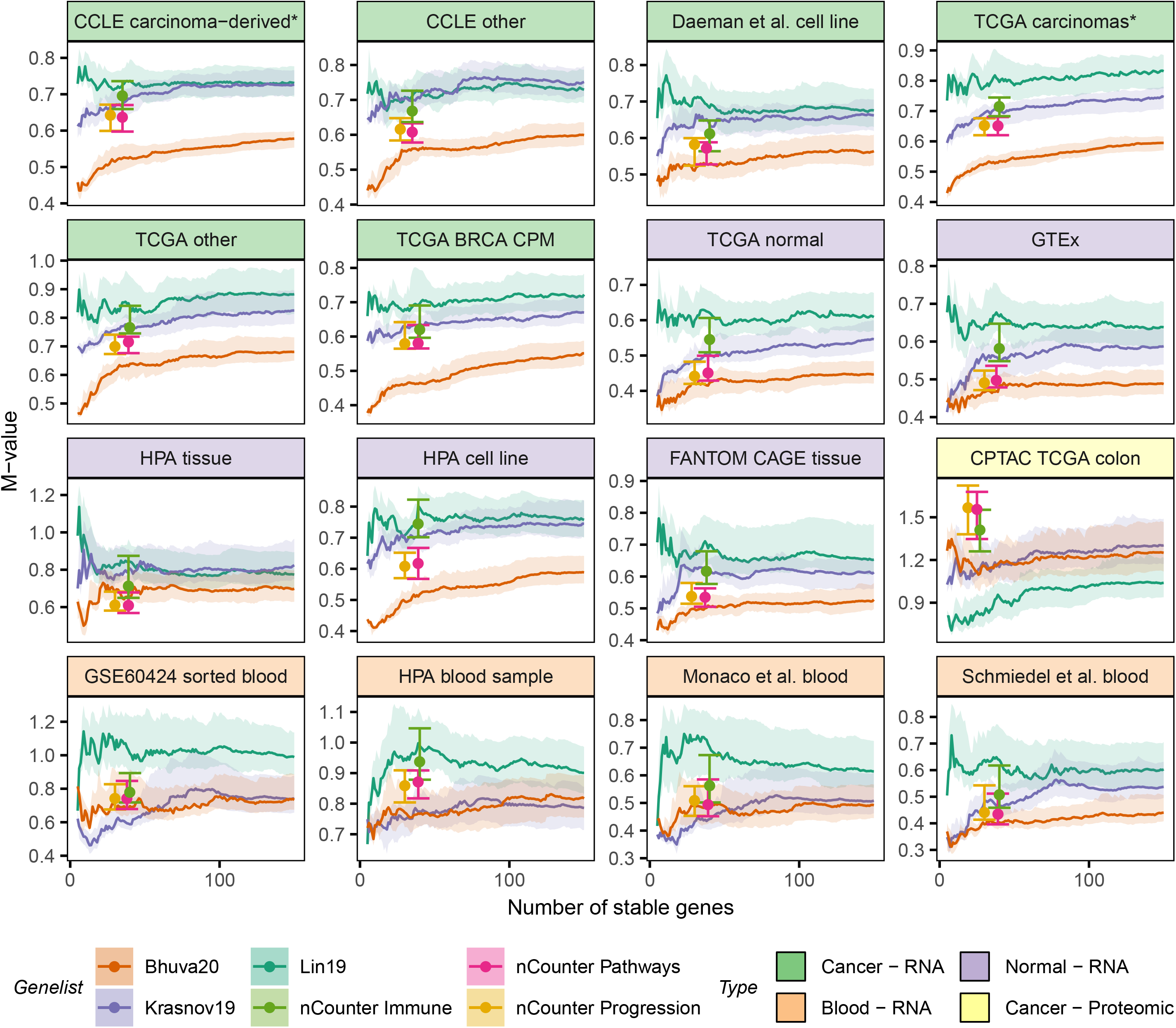
The proposed list is more stable than other lists across most of the 14 datasets used to assess stability. Our prioritisation of stable genes is compared against those from Krasnov et al. (2019) and Lin et al. Stability is compared against fixed lists used in the NanoString nCounter® PanCancer gene panels. Our list results in a better prioritisation of stable genes in cancer patient and cell line datasets. Additionally, our list is more stable in normal samples. Different summarisations of transcriptomic measurments (RPKM, TPM and CPM) do not affect stability. Stability of genes proposed in all lists is not preserved in blood, likely due to the distinct biology of blood. Additionally, our list like most others was identified on solid tissue data, therefore, did not capture stability in blood. Genes proposed by Lin et al. tend to be more stable in proteomic data. This list was proposed for scRNA-seq data which suffers from the same problem of missing values as proteomic data therefore would work well with proteomic measurements.

As shown, our list outperforms others in stability measures across the datasets used for their derivation. Additionally, our list outperforms other lists in all TCGA datasets, including non-carcinomas, normal tissue samples and the breast cancer cohort with gene expression measured in CPM. Interestingly, all lists of stable genes tend to be more stable in normal TCGA samples than in cancer samples as evident by the relatively lower M-values. We note that magnitudes of M-values should not be compared between datasets if data are drawn from different underlying distributions. It is a valid comparison for RPKM-level data from TCGA as they were prepared and processed similarly. Our genes are more stable than other lists across cancer datasets, including CCLE non-carcinoma cell lines and Daemen et al. (34) breast cell lines. Additionally, our genes are relatively more stable across normal tissue and primary cell lines except for the human protein atlas tissue data where the NanoString nCounter® PanCancer Pathways and PanCancer Progression panel genes are more stable compared to our lists of equivalent size. Control genes for the NanoString nCounter® panels were selected by optimising M-values in the GTEx data, yet our lists of equivalent size are more stable in those data. We also showed stability of our list on data generated using a different protocol for measuring transcript expression, CAGE-seq.

The most distinct datasets were the blood RNA-seq and cancer proteomic datasets. None of the lists of stable genes clearly outperform others in blood data. The top 50 stable genes from Krasnov et al. (20) tend to be more stable in blood compared to other lists, but the trend does not hold when more stable genes are considered. Our list outperforms other lists in the blood data generated by Schmiedel et al. (39). Interestingly, control genes used in the NanoString nCounter® PanCancer Immune panel tend to be less stable in blood data than our stable genes and those proposed by Krasnov et al. (20). Finally, the list by Lin et al. (21) is the most stable in the label-free quantification proteomic dataset from the CPTAC project. Overlap analysis between the different lists showed that there was little overlap between our list of stable genes and other lists (Additional file 1: Supplementary figure 2). As such, the list of stable genes we propose is relatively novel. Additionally, the set of reference genes commonly used across many gene expression panels (gathered by Krasnov et al. (20)) have the little overlap with genes identified from data-centric approaches.

### Functional composition of stable genes

Next, we investigated the functional role of genes with stable expression. Since these genes were expressed at stable levels across a wide range of tissues, we suspected they may be involved in essential processes. To test this hypothesis, we used the list of essential genes identified in the DepMap project (40,41) using CRISPR knock-out screens across a variety of cell lines. The DepMap project identified 2164 genes that were essential for survival. We evaluated their occurrence in our list of stable genes and noticed that for lists of any size, approximately half the stable genes were essential for survival. Similarly, approximately 65% of the stable genes proposed by Lin et al. (21), approximately 35% of the genes proposed by Krasnov et al. (20) and approximately 40% of the genes in the NanoString nCounter® panel were essential for survival (see Additional file 1: Supplementary figure 3).

Since stable genes were a mix of essential and non-essential genes, we further characterised our list using gene ontology (GO) enrichment analysis. Our aim was to evaluate enrichment accounting for the prioritisation of stability in our list. We adapted singscore (35) to achieve this and computed stability scores for gene sets derived from gene ontologies and KEGG pathways. Gene sets associated with RNA processing (GO:0006396, GO:0008380), the spliceosome complex (GO:0005681), mRNA metabolic process (GO:0016071) and RNA binding (GO:0003723) were some of the gene sets enriched with stable genes (see Additional file 1: Supplementary figure 4) and had varying proportions of essential genes (50%-75%, see Additional file 2). Similar analysis of KEGG pathways as gene sets revealed the spliceosome pathway (hsa03040) to be the most stable. The epithelial mesenchymal transition gene set from the hallmarks set of MSigDB was the most variable (least stable). Stability scores of all MSigDB gene sets can be further explored using the interactive plot available in Additional File 2.

### Relative ranks of stable genes are preserved across samples

The perfect stable gene would be completely invariant under all conditions. Given two such genes expressed at different abundances, we would observe that across all samples, the gene with higher expression would always be ranked higher than the other gene. This relationship would slowly dissolve as these genes become more variant or the difference between their average expression reduces. As such, given a set of stable genes, their ranks based on gene expression would be consistent so long as the genes were stable. We identified this effect for the highest and lowest expressed genes in our top 5 stable genes, *RBM45* and *HNRNPK* respectively. *RBM45* expression was lower than *HNRNPK* expression for all samples across all 15 RNA-seq datasets. This strong rank preservation is in part attributed to the large difference in expression between the two genes.

Expression-based ranks for genes were computed using the product of ranks meta-analysis approach. We ranked stable genes according to their median expression in the datasets used to identify them (TCGA carcinomas and CCLE carcinoma-derived cell lines). Using the rank of median expression in each dataset as an individual *ranker*, we computed the product of ranks to determine the expression-based rank of each gene. As such, information on the expected ranks based on abundance is added to any discretisation for our list of stable genes. Next, we evaluated rank preservation for stable gene sets of sizes 5 to 30 across all datasets. For each pair of genes within a set of stable genes, we first computed the pairwise rank consistency as the proportion of samples where the order of gene expression matched the expected order. Then, for each gene, we defined the gene-wise rank consistency as the average of its pairwise consistency with all other genes in the set. Like the M-value, the gene-wise rank consistency of a gene is defined relative to other genes in the gene set. Figure 3a shows the pairwise rank consistency measurements for the top 7 stable genes along with their gene-wise rank consistencies in TCGA carcinomas.

**Figure 3.**
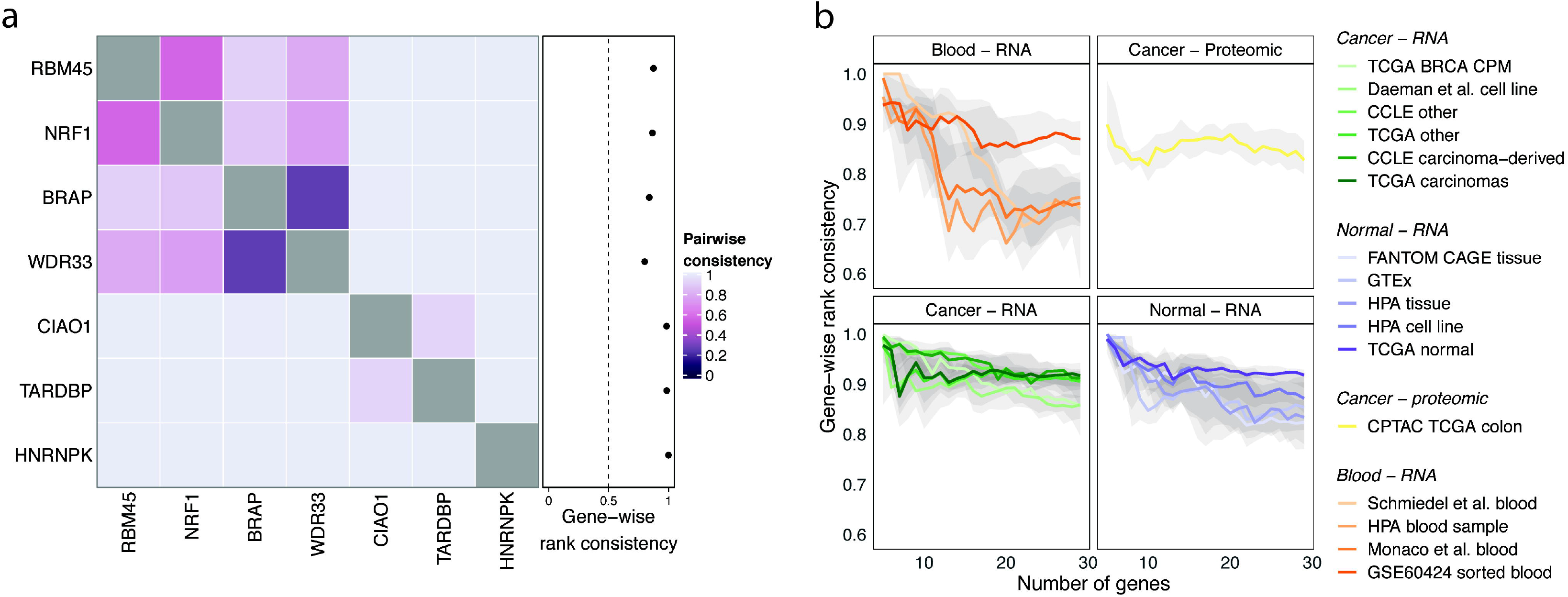
Expression ranks of stable genes are preserved relative to each other. **a)** Pairwise rank consistency measured as the proportion of samples where the expected ranks learned from the training datasets matches the observed rank. The gene-wise rank consistency is then the average of a gene’s pairwise consistencies. The gene-wise consistency of HNRNPK is 1 indicating that its rank relative to the six other stable genes being considered is as expected (higher than the other genes in all samples). **b)** Median gene-wise rank consistencies plot for stable genes from lists of sizes 5 to 30. Bands represent the inter-quartile range. Rank consistency is low in blood datasets due to reduced stability.

We computed gene-wise rank consistencies of genes in stable gene sets of varying sizes across the 14 validation datasets. Figure 3b shows the median of computed gene-wise rank consistencies along with the interquartile range for each set of stable genes. As expected, ranks are preserved in datasets used to derive stable genes and their expected order. Ranks are preserved strongly in cancer datasets and in normal tissue/cell line datasets with a slightly higher preservation in the former. Rank consistency within blood datasets is generally lower compared to all other datasets. Larger sets of stable genes tend to be strongly preserved in TCGA normal samples. Rank preservation is observed in the CPTAC colon proteomic dataset, though not as strongly as with transcriptomic dataset.

### Computing transcriptomic signature scores using a reduced number of measurements

We previously developed a method, singscore, to score individual samples against gene set signatures using transcriptomic data and showed that these scores can assist in assessing the molecular phenotype of tissues and cell lines (35). Though the method has been applied in diverse scenarios in an exploratory context (25,35,36,42,43), the potential for clinical translation is limited by a requirement for transcriptome-wide measurements. Singscore ranked genes based on their transcript abundance, computed the mean of expected up-regulated genes in the case of an up-regulated gene signature, and normalised the mean to produce a signature score. Higher scores indicated concordance with the gene signature. Transcriptome-wide measurements were only required to rank genes in the signature, thereby providing context on how highly/lowly expressed genes in the signature were relative to all other genes.

The rank preservation property of stably expressed genes can be used to calibrate measurements within a sample, thus providing an appropriate context to evaluate the relative expression levels of all other genes. Stable genes allow relative rank estimation of genes without the need for transcriptome-wide measurements. The relative transcriptome-wide rank of any gene can be interpolated given a set of stably expressed genes that are equally spaced on the expression spectrum and that span the entire range of expression values. Our set of genes cover a wide range of the gene expression spectrum and have an approximate uniform distribution (see Figure 1), therefore, may be used to approximate the ranks of other genes without the need for transcriptome-wide measurements. We approximated the unit normalised ranks of genes using a simple approach (see illustration in Additional file 1: Supplementary figure 5). For any given set of stable genes, the rank of a gene was approximated as the number of stable genes with expression values lower than its expression. Unit normalisation was performed by dividing this number by one plus the total number of stable genes in the stable gene set. The signature score of a signature consisting of up-regulated genes was simply the average of their unit normalised ranks. Further normalisation would not be required since ranks were already normalised. A similar procedure would be applied for signatures consisting of down-regulated genes; the only difference being the inversion of ranks (1 – unit normalised rank). Scores using both up- and down-regulated genes were centred around 0 by subtracting the median score (0.5) from them resulting, thus ensuring score were in the range [−0.5, 0.5]. Additionally, signatures composed of both up- and down-regulated genes with unknown direction can be scored using an approach similar to singscore (35).

We compared scores computed using the original singscore with those computed using our new method, *stingscore* (stable singscore), that only requires measurements of genes in the transcriptomic signature and a few stable genes. Samples from the SEQC/MAQCIII project were used to compare scores computed using the different approaches on different measurement platforms (44). The SEQC/MAQCIII project had measurements for four samples: universal human reference RNA sample (UHRR), the human brain reference RNA (HBRR), a mixture containing ¼ UHRR and ¾ HBRR, and a mixture containing ¾ UHRR and ¼ HBRR. HBRR was extracted from multiple brain regions of multiple patients and UHRR was extracted from different tumour tissues of different patients. Measurements of all samples were taken using RNA-seq and RT-qPCR. We scored all samples against a brain-specific neurotransmitter receptor activity gene signature (GO: 0030594) and a cancer-associated cell cycle signature (45). Scores were computed using two approaches: using singscore with transcriptome-wide RNA-seq measurements and using *stingscore* with RT-qPCR measurements of genes in the signatures and the top five stable genes identified in our analysis. Since RT-qPCR measures cycle threshold (C_t_) values, we ranked genes using 1/C_t_. By sampling from the RT-qPCR measurements, we replicate a clinical setting where signatures are evaluated on samples. Scores computed from transcriptome-wide RNA-seq measurements are highly correlated with those from RT-qPCR measurements of signature genes and stable genes (Spearman’s rho > 0.968, Figure 4). Despite the high correlation between scores computed using the different approaches, there is a noticeable yet variable offset in scores. Relevant biology is recapitulated by scores computed using *stingscore*, with HBRR scoring the highest for the brain specific gene signature followed by samples with decreasing amounts of HBRR and finally UHRR having the lowest score. The inverse is noticed with a cell cycle signature which we would expect to be more active in cancers than normal tissue.

**Figure 4.**
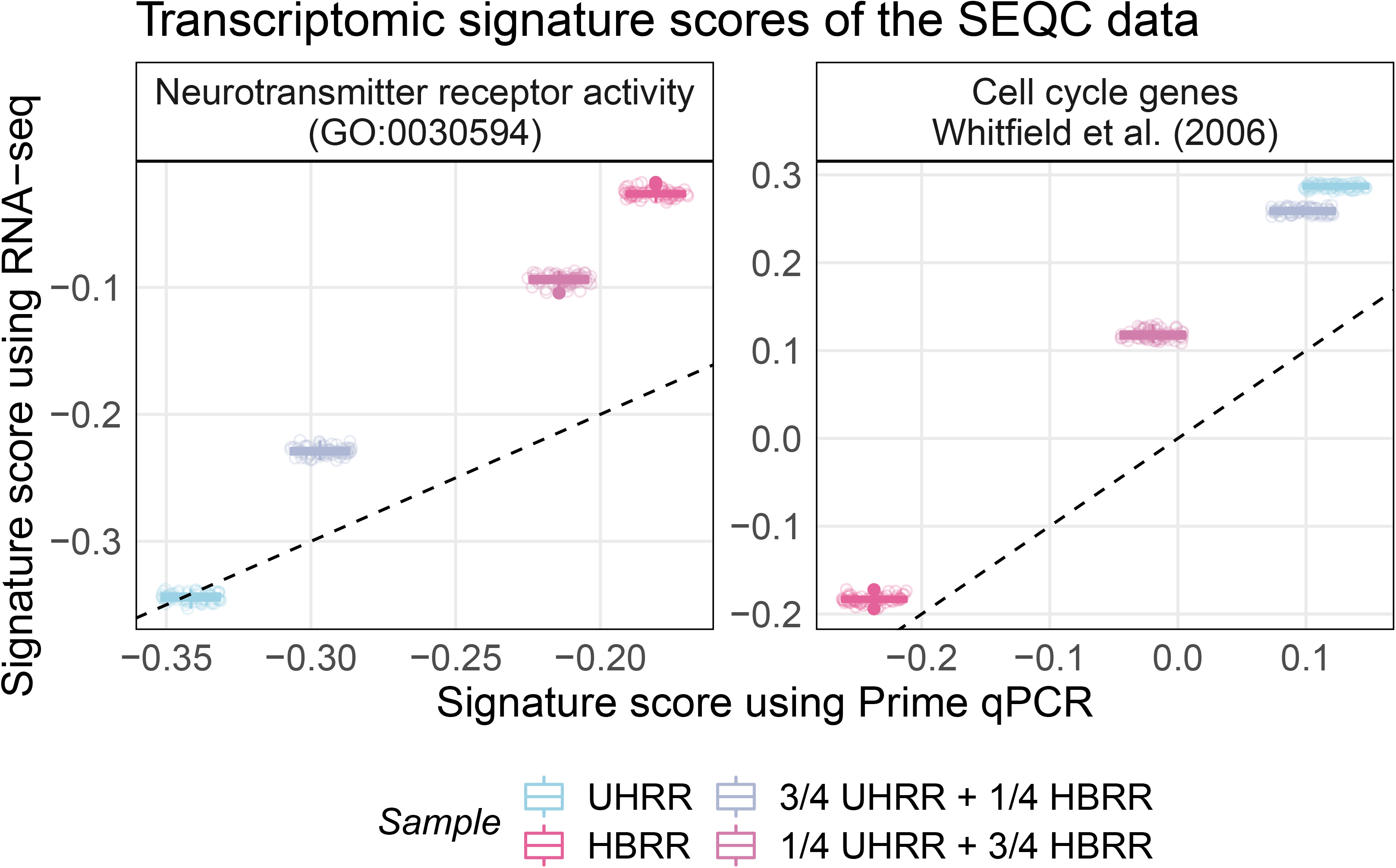
Scores computed from transcriptome-wide RNA-seq measurements are highly correlated with scores computed from a small panel of genes measured using RT-qPCR. The brain RNA sample (HBRR) scores highest for the brain specific signature (neurotransmitter receptor activity) with scores reducing as the proportion of brain RNA reduces. The inverse is observed with a cell cycle gene signature4 which should be more active in cancers. Scores computed using transcriptome-wide RNAseq data are highly correlated (ρ > 0.95) with those computed using RT-qPCR measurements of signature genes and 5 stable genes. RNA-seq data contains multiple observations of the same sample therefore points are jittered along the x-axis for visualisation purposes.

## Discussion

Genes with stable expression have frequently been used for data normalisation including correction of batch effects (9,10). In this study, we derive a new set of stable genes for application in cancer transcriptomic analyses and demonstrate their additional use for targeted molecular phenotyping beyond data normalisation. Targeted molecular phenotyping is particularly useful in the analysis of archival tissue where rich clinical information is present but the preservation techniques limit extraction of high-quality RNA and consequently RNA sequencing. These samples are suitable for other smaller scale measurement platforms such as NanoString® and RT-qPCR. Thus, exploration of strategies such as *stingscore* that move from reliance on a whole transcriptome of measurements to the measurement of tens or hundreds of transcripts will support the analysis of these valuable and vast historical collections of tissues.

We formulated the identification of stable genes as a meta-analysis problem and used the product of ranks approach to combine multiple stability metrics across two datasets. Our approach provides flexibility in adding/removing variability metrics and datasets. Since the product of ranks requires all rankers (the test metrics) to provide a complete ranking, we had to discard genes that were not measured and consequently not ranked in either dataset. While this loss was relatively small for integration of two datasets, it would be more pronounced if multiple datasets were used. Other methods that allow rankers to provide partial rankings might be used (46), but in the specific application of identifying a general set of stably expressed genes, it makes sense to limit our analysis to genes that can be measured reliably.

Our set of stable genes covered a wide range of expression values (see Figure 1) even though this property was not selected for explicitly. This is likely a result of using multiple measures of variation which reduced biases resulting from the mean-variance relationship often observed in RNA-seq data (47). A wide range of expression values is desirable for normalisation, as such genes capture variation at the tails of the gene expression spectrum. A wide range is also important for the evaluation of gene expression signatures that capture down regulated genes, where we expect low abundance. Our selection of stable genes also minimises between-group differences by minimising the F-statistic from a one-way ANOVA test on groups. Stable genes do not always have the smallest F-statistics, but this is expected. Small between-group differences for highly variable genes are less significant and will produce smaller F statistics compared to the same magnitude of differences for less variable genes. While stability of genes is assessed relative to other genes, this metric still holds value in our analysis.

We validated stability of our genes across multiple independent datasets. To our knowledge, this was the first study to evaluate stability of genes across such a large and diverse set of data. Our evaluation of stability was performed across 14 datasets (with approximately 13000 samples) representing different biology (cancer, normal, and blood), collected by different consortia using different preparation protocols, measured with different instruments, processed using different pipelines and summarised using different metrics. Our set of stable genes were equally or more stable than other lists of stable genes in cancer and normal tissue and cell line datasets as shown by the smaller M-values (see Figure 2). Stability of all lists was poor and inconsistent in blood datasets, likely due to the genes being originally identified as stable in samples derived largely from solid tissue which is biologically very distinct from haematological and lymphoid tissues and cells. Genes with stable expression in blood could easily be identified by using our approach and appropriate data. An interesting observation was the stability of genes identified by Lin et al. (21) in label-free protein quantification data. Single-cell transcriptomic datasets suffer from the same problem of missing values as label free quantification, therefore genes determined to be stable in one will likely perform well in the other. Our list of stable genes and by extension other lists do not necessarily have poor stability in proteomic data, but they may be more difficult to measure. Stability is a prerequisite to rank preservation as evident from the low rank preservation in blood and proteomic data and high rank preservation in normal tissue.

Investigation into the functional roles of our proposed stable genes revealed that genes essential for cell survival tend to exhibit stable expression, however this is not universal. Further exploration of gene ontologies and other gene expression signatures from MSigDB revealed molecular processes involving RNA processing to be enriched in stably expressed genes. This observation indicates that these processes essential for cell survival are finely regulated and little room for error exists. Additionally, we showed that context/tissue-specific processes such as epithelial-mesenchymal transition were enriched with highly variable genes, as expected given the diverse changes associated with this phenotypic program (43). We used singscore (35) with stability ranks to enable this analysis, demonstrating that singscore can be used to assess enrichment using any ranked data.

Using the rank preservation property of stable genes, we developed a new molecular phenotyping method, *stingscore,* based on our original method (35). To date, this is the only approach capable of computing signature scores for single samples using a reduced set of transcriptomic measurements, such those obtained in a targeted study or RT-qPCR panel. We demonstrated that signature scores computed using the two methods and measurements are only strongly associated if they were computed using biologically meaningful gene signatures (see Figure 4). Such signatures possess the power to discriminate samples therefore scores are correlated between approaches. In contrast, non-relevant signatures capture noise in the context of the biological problem being analysed. The choice of signatures used to analyse any biological problem should be motivated by prior knowledge of the biological system being analysed. Though scores generated using singscore and *stingscore* are correlated, they are not equivalent and there is generally an offset (see Figure 4). Ranked sample order however is preserved. This variation could be addressed by identifying other stable genes that could be used to calibrate scores. For instance, we could identify a stable gene set that represent the median score observed across all samples and adjust the score of each sample against the stable gene set score such that positive scores indicate concordance with a gene signature and vice versa.

Our results demonstrate the potential for stable genes in clinical translation of biologically relevant gene sets through single sample transcriptomic gene signature scoring with a reduced panel of target genes. More sophisticated calibration methods and scoring methods capable of application to single samples, or small numbers of samples often resulting from clinical research, will be enabled by the ideas and methods presented in this work.

## Conclusion

Molecular profiling at the individual patient level is becoming increasingly useful in the clinic despite the lack of translation of such approaches. A wealth of molecular gene signatures such as those in the Molecular signatures database (MSigDB), remain unexplored in the clinic because of the requirement of whole transcriptome measurements imposed by most computational approaches that limit deployment in the context of regular pathology testing. In this study, we propose a novel cost-effective panel-based approach using stably expressed genes to assess transcriptomic gene signatures for individual patients in the clinic based on measurement of a substantially reduced number of genes (around two to three orders of magnitude fewer than whole transcriptome scale measurement). Since stable genes are used in this approach, no additional genes are required for data normalisation thus further saving costs of panel-based tests. This method will facilitate the adoption of transcriptomic gene signatures analysis in a clinical context, thus allowing molecular profiling of a patient’s disease, along with assessment of diagnostic/prognostic gene signatures, and assessment of signatures predictive of response to therapies.

## Supporting information

Additional file 2 (HTML)

## Declarations

### Availability of data and material

Methods developed in this study and the complete list of stable genes in human carcinomas are available in the R/Bioconductor package `singscore` (v1.8.0).

### Supplementary data

Additional file 1: Supplementary methods and figures. This file contains additional results generated to support the findings in this study. (PDF 369 KB)

Additional file 2: Interactive figure. This file contains an interactive figure where the stability of gene signatures in the molecular signature database (MSigDB) is plot. Stability scores for gene signatures are plot against their dispersion estimates. (HTML 6.44 MB)

Additional file 3: R code. This file contains the functions used to compute stability measures for datasets and subsequently extract stably expressed genes. (R 5 KB)

## Acknowledgements

The Genotype-Tissue Expression (GTEx) Project was supported by the Common Fund of the Office of the Director of the National Institutes of Health, and by NCI, NHGRI, NHLBI, NIDA, NIMH, and NINDS. The data used for the analyses described in this manuscript were obtained from the GTEx Portal on 13/08/2019.

## Funding

This work was supported by National Health and Medical Research Council funding (Project Grant 1128609 to MJD, Project Grant 1147528 & 1165208 to JC), Cancer Council Victoria funding (Project grant 1187825 to MJD), and National Breast Cancer Foundation and Cure Brain Cancer Foundation funding (Project Grant CBCNBCF-19-009 to MJD). MJD is the recipient of the Betty Smyth Centenary Fellowship in Bioinformatics. DDB was supported by the Melbourne Research Scholarship (MRS). This study was made possible through Victorian State Government Operational Infrastructure Support and Australian Government NHMRC Independent Research Institute Infrastructure Support scheme.

## Competing interests

The authors declare no competing interests.

## Authors’ contributions

DDB and MJD conceived and designed this study. DDB developed the methodology and software, and performed the computational analyses described. JC and MJD supervised this study. All authors wrote and approved the manuscript.

## Notes

https://doi.org/doi:10.18129/B9.bioc.singscore

### Competing Interest Statement

The authors have declared no competing interest.

