## Additional file 1 (PDF) for "Stable gene expression for normalisation and single-sample scoring"

### Supplementary figures

| | TCGA carcinomas (ranks) | | | | CCLE carcinoma-derived (ranks) | | | | | |
|---|---|---|---|---|---|---|---|---|---|---|
| | MAD | ENT | OSS | F | MAD | ENT | OSS | F | PR | Rank |
| g1 | 1 | 1 | 2 | 3 | 1 | 2 | 1 | 4 | 48 | 1 |
| g2 | 3 | 2 | 1 | 2 | - | - | - | - | - | - |
| g3 | 2 | 3 | 4 | 4 | 3 | 1 | 2 | 3 | 1728 | 2 |
| g4 | 5 | 5 | 5 | 1 | 2 | 3 | 4 | 1 | 3000 | 3 |
| g5 | 4 | 4 | 3 | 5 | 4 | 4 | 3 | 2 | 23040 | 4 |

***Supplementary figure 1: Product of ranks approach to stable genes prioritisation.***

*The median absolute deviation (MAD), Shannon's entropy (ENT), outlier sum statistic (OSS) and the F-statistic from an ANOVA analysis on the groups is performed in each of the two datasets. Genes 1 to 5 are then ranked for each statistic in each dataset (rank along columns). The product of these ranks for each gene across all datasets and statistics produces the product of ranks statistic (PR). Genes can then be ranked and prioritised using this statistic. Genes missing in any dataset are discarded as their complete product of ranks cannot be computed.*

##### Overlap analysis

| | Bhuva20 (top 5) | Bhuva20 (top 30) | Bhuva20 (top 150) | nCounter PanCancer Pathways | nCounter PanCancer Immune | nCounter PanCancer Progression | Reference | Krasnov19 (top 150) | Lin19 (top 150) | LINCS1000 | Eisenberg 2013 |
|---|---|---|---|---|---|---|---|---|---|---|---|
| Bhuva20 (top 5) | 5 | 5 | 5 | 1 | 0 | 0 | 0 | 1 | 2 | 0 | 3 |
| Bhuva20 (top 30) | 5 | 30 | 30 | 4 | 3 | 3 | 0 | 3 | 5 | 3 | 25 |
| Bhuva20 (top 150) | 5 | 30 | 150 | 11 | 9 | 8 | 1 | 12 | 13 | 12 | 114 |
| nCounter PanCancer Pathways | 1 | 4 | 11 | 40 | 30 | 30 | 0 | 2 | 1 | 4 | 24 |
| nCounter PanCancer Immune | 0 | 3 | 9 | 30 | 40 | 30 | 8 | 3 | 1 | 7 | 22 |
| nCounter PanCancer Progression | 0 | 3 | 8 | 30 | 30 | 30 | 0 | 2 | 1 | 3 | 18 |
| Reference | 0 | 0 | 1 | 0 | 8 | 0 | 32 | 0 | 2 | 7 | 14 |
| Krasnov19 (top 150) | 1 | 3 | 12 | 2 | 3 | 2 | 0 | 150 | 1 | 22 | 102 |
| Lin19 (top 150) | 2 | 5 | 13 | 1 | 1 | 1 | 2 | 1 | 150 | 9 | 110 |
| LINCS1000 | 0 | 3 | 12 | 4 | 7 | 3 | 7 | 22 | 9 | 978 | 338 |
| Eisenberg 2013 | 3 | 25 | 114 | 24 | 22 | 18 | 14 | 102 | 110 | 338 | 3803 |

***Supplementary figure 2: Proposed stable genes have little overlap with previously identified sets.*** *The top 5, 30 and 150 stable genes from our analysis are compared against stable genes from the NanoString® nCounter pancancer analysis panels, a set of reference genes commonly used in gene expression panels (gathered by Krasnov et al. (2019), top 150 stable genes identified by Krasnov et al. (2019), top 150 stable genes identified by Lin et al. (2019), genes identified as housekeeping genes by Eisenberg et al. (2013) and genes in the LINCS1000 panel that are thought to represent the entire transcriptome. Little overlap exists*

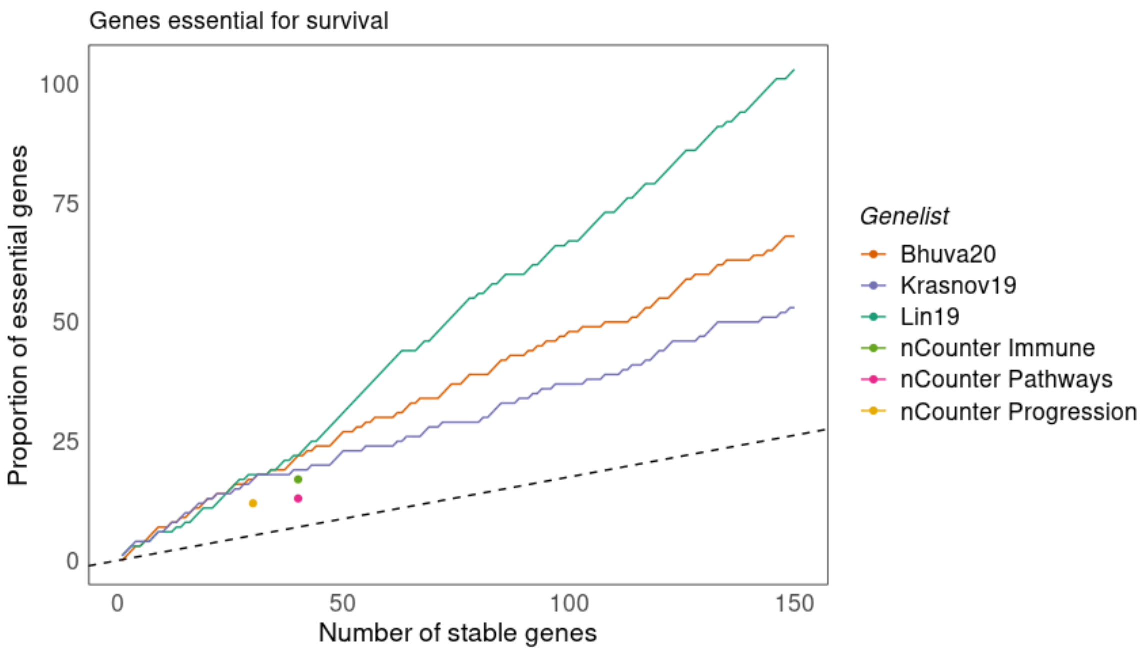


***Supplementary figure 3: Genes essential for cell survival are enriched in sets of stable genes.***

*Genes essential for survival as identified in the DepMap project are overlapped with stable gene sets of varying sizes. Sets of different size were tested and the number of essential genes in them are plot. The expected number of essential genes in random sets is shown by the dashed line. All sets are composed of essential genes and are enriched in them.*

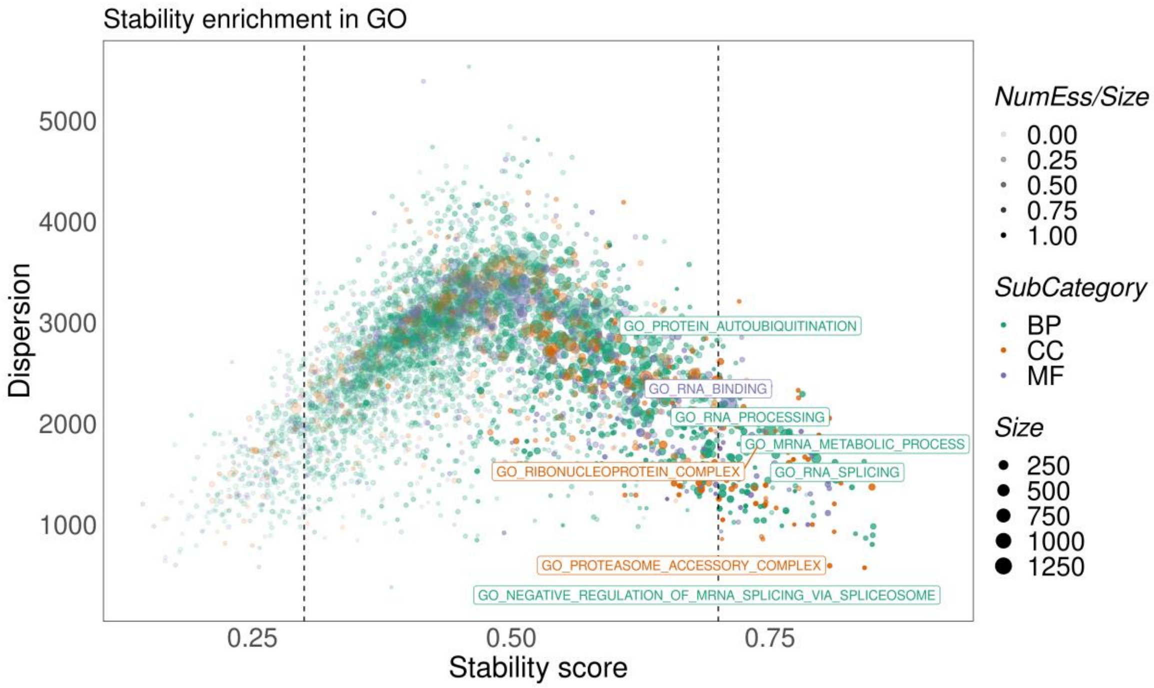


***Supplementary figure 4: Stability analysis of gene ontology derived gene sets.***

*Gene sets formed from gene ontologies are scored for stability and annotated by the proportion of essential survival genes present in them (transparency). Gene sets with scores close to 1 are enriched in stable genes. Gene sets with many essential genes tend to be composed of stable genes but this is not always the case (scores between 0.5 and 0.7). Gene sets associated with RNA processing, RNA splicing, RNA binding and the proteasome are enriched in stably expressed genes.*

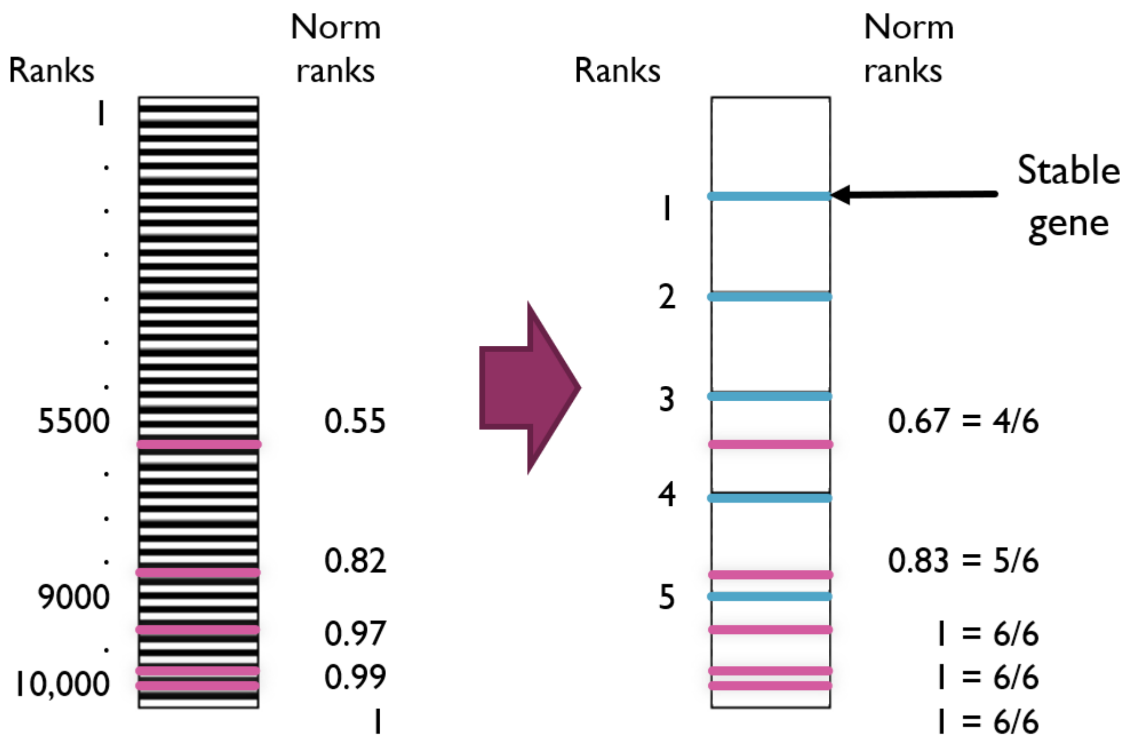


***Supplementary figure 5: Rank approximation using stable genes.***

*The unit normalised expression-based rank of genes (e.g. purple) can be approximated with only a few stable genes (blue). The rank of gene of interest (purple gene) is computed as the number of stable genes its expression is larger than (e.g. 3 for the gene with the lowest expression) divided by 1 + total number of stable genes used. As can be seen above, unit normalised ranks for the genes using the entire transcriptome (left) are well approximated by using just 5 stable genes (right). Approximated ranks can then be used to compute signature scores where genes in the signature are the genes of interest (purple).*
