## Additional file 3 (R code) for "Stable gene expression for normalisation and single-sample scoring": additional_file_3.R

```
library(entropy)
library(foreach)

#' Compute stability metrics for a single gene expression profile
#'
#' @param e a numeric vector, containing the expression profile of a single gene
#' @param annot a factor, containing the annotation for each sample
#' @param full_erange a numeric, (1 x 2) containing the range of expression for
#'   the entire dataset
#'
#' @return A vector containing summary metrics for the expression profile.
#'   Stable genes have lower values for each metric (excluding avg & gene)
#'
stabilityMetricsGene <- function(e, annot, full_erange) {
  #convert annotation to factors
  smetrics = list(
    'avg' = median(e),
    'mad' = mad(e),
    'entropy' = entropy::entropy(entropy::discretize(e, floor(sqrt(length(e))), r = full_erange)),
    'class_diff' = summary(aov(e ~ annot))[[1]][1, 4]
  )
  
  #calculation for outliers
  e = e - smetrics$avg
  qts = quantile(e, c(0.25, 0.75))
  outliers = e[e > qts[2] * 2 - qts[1] | e < qts[1] * 2 - qts[2]] #(q0.75 + IQR)
  smetrics = c(smetrics, list(
    'outlier_sum' = sum(abs(outliers)),
    'outlier_num' = length(outliers)
  ))
  
  return(smetrics)
}

#' Compute stability metrics for all genes in a dataset
#'
#' @param edata a matrix, of expression values. Genes are along the rows and
#'   samples along the columns
#' @param annot a factor, containing the annotation for each sample
#'
#' @return A data.frame containing summary metrics for the expression profile.
#'   Stable genes have lower values for each metric (excluding avg & gene).
#'   Genes are sorted based on the stability metric (prod_rank)
#'
computeStabilityMetrics <- function(edata, annot) {
  edata = edata[sort(rownames(edata)), ]
  annot = as.factor(as.character(annot))
  erange = range(edata)
  metricdf = foreach(
    e = iterators::iter(edata, by = 'row'),
    .combine = rbind,
    .inorder = FALSE,
    .packages = c('entropy')
  ) %dopar% {
    metrics = stabilityMetricsGene(e[1, ], annot, erange)
    metrics$gene = rownames(e)
    return(data.frame(metrics, row.names = metrics$gene))
  }
  
  #select measures to use to compute the product of ranks
  prod_cols = c('mad', 'entropy', 'class_diff', 'outlier_sum')
  
  #compute the product of ranks and sort accordingly
  metricdf$log_pr = apply(log(apply(metricdf[, prod_cols], 2, rank)), 1, sum)
  metricdf = metricdf[order(metricdf$log_pr), ]
  
  return(metricdf)
}

#' @title Get a list of stably expressed genes
#'
#' @description Get a list of genes that are stably expressed in given stability
#'   measures of a dataset
#'
#' @param n numeric, number of stable genes to retrieve
#' @param ... metric dataframes returned from the `computeStabilityMetrics`
#'   function
#' @param ordAvg logical, if TRUE, genes are ordered based on their median
#'   expression across datasets
#'
#' @return a character vector with gene IDs sorted by their expected expression
#'   levels in the requested tissue
#'   
getStableGenes <- function(n, ..., ordAvg = TRUE) {
  metricdfs = list(...)
  common_genes = metricdfs[[1]]$gene
  for (i in 2:length(metricdfs))
    common_genes = intersect(common_genes, metricdfs[[i]]$gene)
  
  #subset to common genes
  metricdfs = lapply(metricdfs, function(x) x[common_genes, ])
  
  #re-rank
  log_pr = rowSums(data.frame(lapply(metricdfs, function(x) log(rank(x$log_pr)))))
  common_genes = common_genes[order(log_pr)]
  
  #select top N genes
  stopifnot(n <= length(common_genes))
  genes = common_genes[1:n]
  
  if(ordAvg) {
    avgranks = rowSums(data.frame(lapply(metricdfs, function(x) log(rank(x[genes, 'avg'])))))
    genes = genes[order(avgranks)]
  }
  
  return(genes)
}

#----example----
set.seed(36)

#create data
ds1 = matrix(rnorm(100), 10, 10)
ds2 = matrix(rnorm(100), 10, 10)
group1 = group2 = rep(1:2, each = 5)
#8 common genes
rownames(ds1) = LETTERS[1:11][-1]
rownames(ds2) = LETTERS[1:11][-3]

#compute metrics
metricdf1 = computeStabilityMetrics(ds1, group1)
metricdf2 = computeStabilityMetrics(ds2, group2)
getStableGenes(5, metricdf1, metricdf2) #F K J I G (for R > 3.6)
getStableGenes(5, metricdf1, metricdf2, ordAvg = FALSE) #J F G K I (for R > 3.6)
```
